# Characterising differential gene expression and alternative splicing in a sex reversing skink, *Bassiana duperreyi*

**DOI:** 10.64898/2026.06.15.731768

**Authors:** Benjamin J. Hanrahan, J King Chang, Duminda S.B. Dissanayake, Nicholas C. Lister, Arthur Georges, Paul D. Waters

**Author notes:** Correspondence: Paul D. Waters.

## Abstract

In some reptiles, genetic and environmental sex determination interact whereby extreme incubation temperatures override genetic sex determination (GSD) to produce sex-reversed individuals. In one lizard with temperature-influenced GSD, the central bearded dragon, intron retention in the histone-modifier genes *Kdm6b* and *Jarid2* has been implicated as a candidate signal linking temperature to sex. Equivalent intron retention is also present in two species with temperature-dependent sex determination, the red-eared slider turtle and the American alligator. The eastern three-lined skink, *Bassiana duperreyi*, represents another lizard with temperature induced sex reversal. It has an XY sex determination system in which low temperature incubation causes sex reversal of XX embryos to produce phenotypic males.

In this study, we performed splice-aware analysis of RNA sequencing from hatchling brains of the three-lined skink. We investigated differences in alternative splicing and gene expression between the three sex conditions: XY males (XYm), XX females (XXf), and sex-reversed XX males (XXm). Sex reversal specific intron retention was observed in the gene, *Ttll7*, which only occurred in XXm and not in XYm or XXf. Intron retention in *Ttll7* could alter the function of the encoded protein, a tubulin polyglutamylase, but its effect on sex reversal here is unknown.

In addition, intron retention in the histone-modifier genes *Jarid2* and *Kdm6b* occurred in all conditions. The presence of *Kdm6b* and *Jarid2* intron retention in all sex conditions suggests that the pattern of intron retention in sex reversal in the eastern three-lined skink is distinct compared to the bearded dragon. We conclude that a different molecular pathway for sex reversal is induced in the three-lined skink, the details of which remain elusive.

## Introduction

Sex determination occurs at a developmental point at which bipotential gonads start the process of developing into a testis or an ovary. In vertebrates this is generally either a genetic cue (genetic sex determination – GSD) or an environmental signal such as temperature (temperature-dependent sex determination – TSD). In GSD systems, sex chromosomes have a different complement between males and females, which ultimately determines sex. In contrast, TSD systems rely on different incubation temperatures during a thermosensitive period to affect clutch sex ratios (Bull 1980; Bull 1983). In some reptile species these genes and environment interact, whereby an underlying GSD signal can be overridden by incubation temperature extremes during development to produce sex-reversed individuals (Shine et al. 2002; Quinn et al. 2007; Radder et al. 2008). This occurs in both the laboratory and wild populations (Holleley et al. 2015; Dissanayake et al. 2021a; Dissanayake et al. 2021b). The interaction of temperature and genetics provides an opportunity to study how the thermal signal at extremes is captured during sex reversal and in TSD. Despite recent research, the signal that transmits the temperature signal to influence sex determination remains unknown (Yatsu et al. 2016; Ge et al. 2018; Whiteley et al. 2022).

Two lizards with temperature induced sex reversal are the central bearded dragon, *Pogona vitticeps* (Quinn et al., 2007; Holleley et al., 2015) and the eastern three-lined skink, *Bassiana duperreyi* (Shine et al. 2002; Radder et al. 2008; Dissanayake et al. 2021b). Work on the central bearded dragon has identified alternative splicing and intron retention (IR) as mechanisms of interest when investigating temperature-induced sex reversal (Deveson et al. 2017). Specifically, two genes that function in histone modification have been identified as having temperature-induced intron retention in the bearded dragon as well as two distantly related TSD species, alligator and turtle. These genes are *Jarid2* (Jumonji and AT-rich interaction domain containing 2) and *Kdm6b* (lysine demethylase 6B) (Deveson et al. 2017; Whiteley et al. 2022).

The eastern three-lined skink, *B. duperreyi*, is an oviparous scincid lizard with an XY sex determination system (Donnellan 1985; Cogger 2014; Matsubara et al. 2016). At low temperatures during incubation in the egg, *B. duperreyi* displays female to male sex reversal causing the development of XXm males (Quinn et al. 2009). Unlike *P. vitticeps, B. duperreyi* has differentiated sex chromosomes with the Y chromosome being gene poor compared to the X, and partial dosage compensation of the X in males (Hanrahan et al. 2026). There is little known about the sex determination of *B. duperreyi* or what genes may be involved in sex determination or sex reversal.

Here we present differential gene expression and differential intron retention analyses in the eastern three-lined skink, *Bassiana duperreyi* (Gray, 1838). Gene expression analysis of hatchling brains showed that sex-reversed XXm males are more like XXf females than XYm males in their transcriptomic profile, suggesting the trigger for sex reversal is subtle. Intron retention analysis identified a gene encoding a microtubule polyglutamylase, *Ttll7*, as having differential intron retention in sex-reversed XXm males compared to XXf females and XYm males. This may indicate altered brain function in sex-reversed individuals. We also investigated two histone modifier genes, *Kdm6b* and *Jarid2*, which are implicated in other temperature sex reversal systems and found that intron retention occurs in all sexes in *B. duperreyi*. This indicates that higher temperatures are needed for the complete splicing of the retained introns in these two genes and suggests they may not be involved in sex reversal of *B. duperreyi*.

## Results

### Differential gene expression between sex conditions

We performed differential gene expression analysis on brain tissue samples of *B. duperreyi* hatchlings comparing the three sexes: XXf, XXm and XYm. Multidimensional scaling (MDS) of global gene expression showed XY males separate from both XX females and XX males (Figure 1A). This difference was not exclusive to a single axis. XX males and XX females were not separated in the MDS plot suggesting no substantial global expression change associated with sex reversal. A heatmap of the top 50 most expressed genes does not show any distinct separation of samples into sex condition (Figure 1B). Notably two genes which have a role in chromatin accessibility: jumonji and AT-rich interaction domain containing 2 (*Jarid2*) and lysine demethylase 6B (*Kdm6b*, also called *Jmjd3*), were both in the top 50 highly expressed genes.

**Figure 1.**
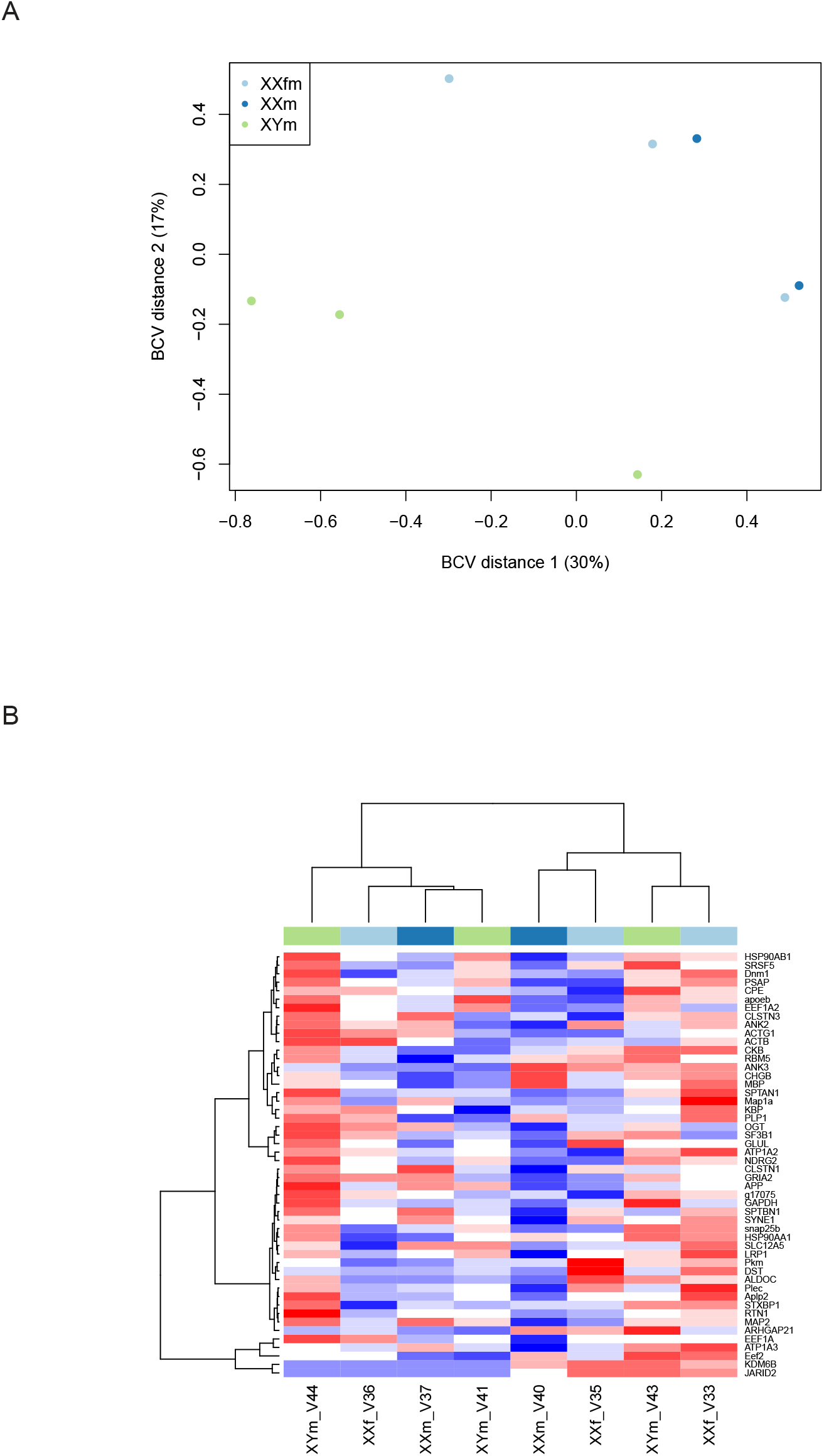
Gene expression overview of the three sexes of *B. duperreyi*. Gene expression analysis of *B. duperreyi* brain samples. **A)** multidimensional scaling (MDS) plot of normalised gene expression shows separation of XYm samples from XXm and XXf samples. Distances correspond to the biological coefficient of variation (BCV) between each pair of samples. **B)** Heatmap of the 50 most highly expressed genes by median expression. Each tile represents the normalised gene expression (Z-score), which was calculated by subtracting the mean expression value of a gene across all samples from the sample specific expression value and divided by the standard deviation of the mean expression of each gene. Distance was calculated using Manhattan distance and hierarchical clustering was calculated using Ward’s method. Samples do not group by sex condition shown by colour above each column, XXf female (light blue), XXm male (dark blue) and XYm male (green).

We were interested in the comparison of sex-reversed XXm to XXf as these conditions share the same genotype, only differing in phenotypic sex because of incubation temperature. However, few genes were found to be differentially expressed (FDR; p < 0.05) between XXf and XXm samples (only 4 total) (Table S1). This suggests that there is not a global expression difference between sex-reversed XXm and XXf that can be linked to sex reversal in our dataset. In addition, none of the four genes: *Cep295, Cabin1, Megf6* or *g11335* (identified as *Rcvrn*, see Table S2), have a known function related to sex determination.

A higher number of genes were found to be differentially expressed (FDR; p < 0.05) in the comparison between XYm and the two XX sex phenotypes (35 for XXf vs XYm and 32 for XXm vs XYm, Table S1). Gene ontology of differentially expressed genes between XYm males and both XX sex conditions identified terms related to brain development and central nervous system related functions such as retina development (see Table S3, S4). None of these genes had functions related to sex determination and were not investigated further.

### Intron retention in sex-reversed XXm males

We next investigated whether sex-reversed *B. duperreyi* displayed differential gene transcript regulation. To do this we performed intron retention (IR) analysis on our samples as differential IR was observed between sexes in another lizard species with sex reversal (the central bearded dragon), as well as two TSD reptiles (the red-eared slider turtle and the American alligator) (Deveson et al. 2017; Whiteley et al. 2022). An analysis of IR events across the brain transcriptome in *B. duperreyi* identified a single gene, tubulin tyrosine ligase-like 7 (*Ttll7*), that contained a differentially retained intron between XXm males compared to both XXf females and XYm males (Table S5). Intron retention of intron 16 in *Ttll7* occurred only in XXm males with an average IR ratio of 36% with read coverage across most of the intron (Figure 2 A-B). *Ttll7* encodes a beta-tubulin polyglutamylase that exhibits brain specific expression (Ikegami et al. 2006). It is not known to have a role in sex determination. Retention of intron 16 would introduce a premature stop codon to the mRNA sequence of *Ttll7* (Figure 2C).

**Figure 2.**
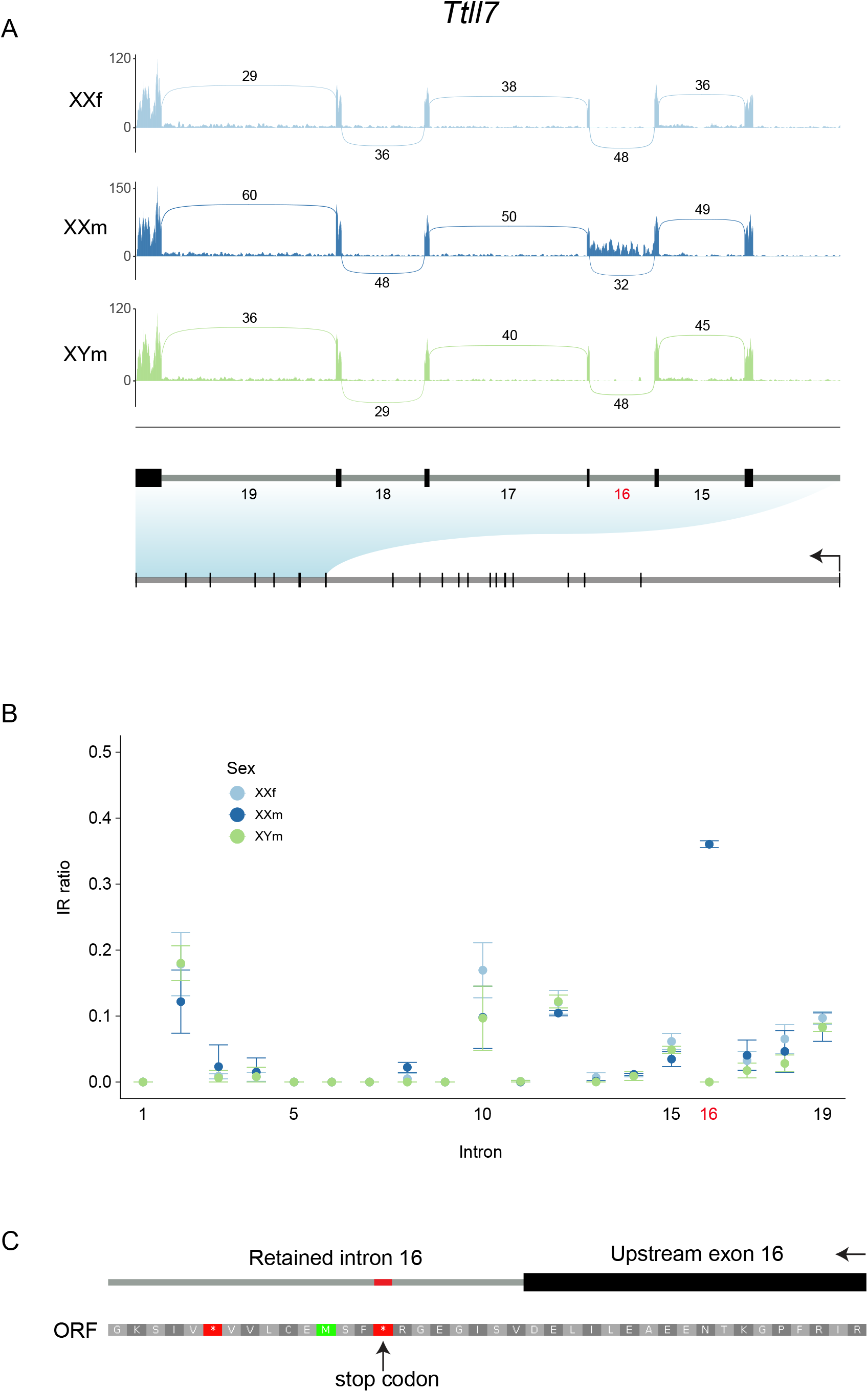
*Ttll7* intron 16 is differentially retained in sex-reversed XXm males. **A)** RNA-seq coverage of *Ttll7* exons 15-20. Tracks show the mean coverage for each sex condition, XXf (light blue, n=3), XXm (dark blue, n=2) and XYm (green, n=3). The mean number of reads spanning exon junctions are shown as lines across the intron above or below the track for each condition. Exon and intron positions are shown below the sashimi plot, including whole gene context. **B)** Intron retention ratio for each intron of *Ttll7* as reported by IRFinder. Points represent the mean for each sex condition, XXf (light blue), XXm (dark blue) and XYm (green) and error bars are standard error. **C)** Exon-intron boundary for exon/intron 16 and translated protein sequence from the canonical open reading frame of *Ttll7*. Stop codons introduced in the retained intron are shown in red.

If translated, the intron-retained isoform of *Ttll7* would result in a truncated protein product that is missing a portion of the C-terminal end of the protein (Figure 3). This truncation would not affect the catalytic tubulin tyrosine ligase-like domain or microtubule binding domain (MTBD) of TTLL7 (Figure 3A). Protein alignment of predicted models showed confident alignment of the area corresponding to the catalytic domain and MTBD between TTLL7 and TTLL7-IR but poor alignment toward the C-terminal end of the protein (Figure 3B). AlphaFold prediction of the canonical TTLL7 resolves the tertiary structure of the C-terminal end with high confidence (low position error). In comparison this section is missing in the TTLL7-IR prediction due to intron retention (Figure 3C).

**Figure 3.**
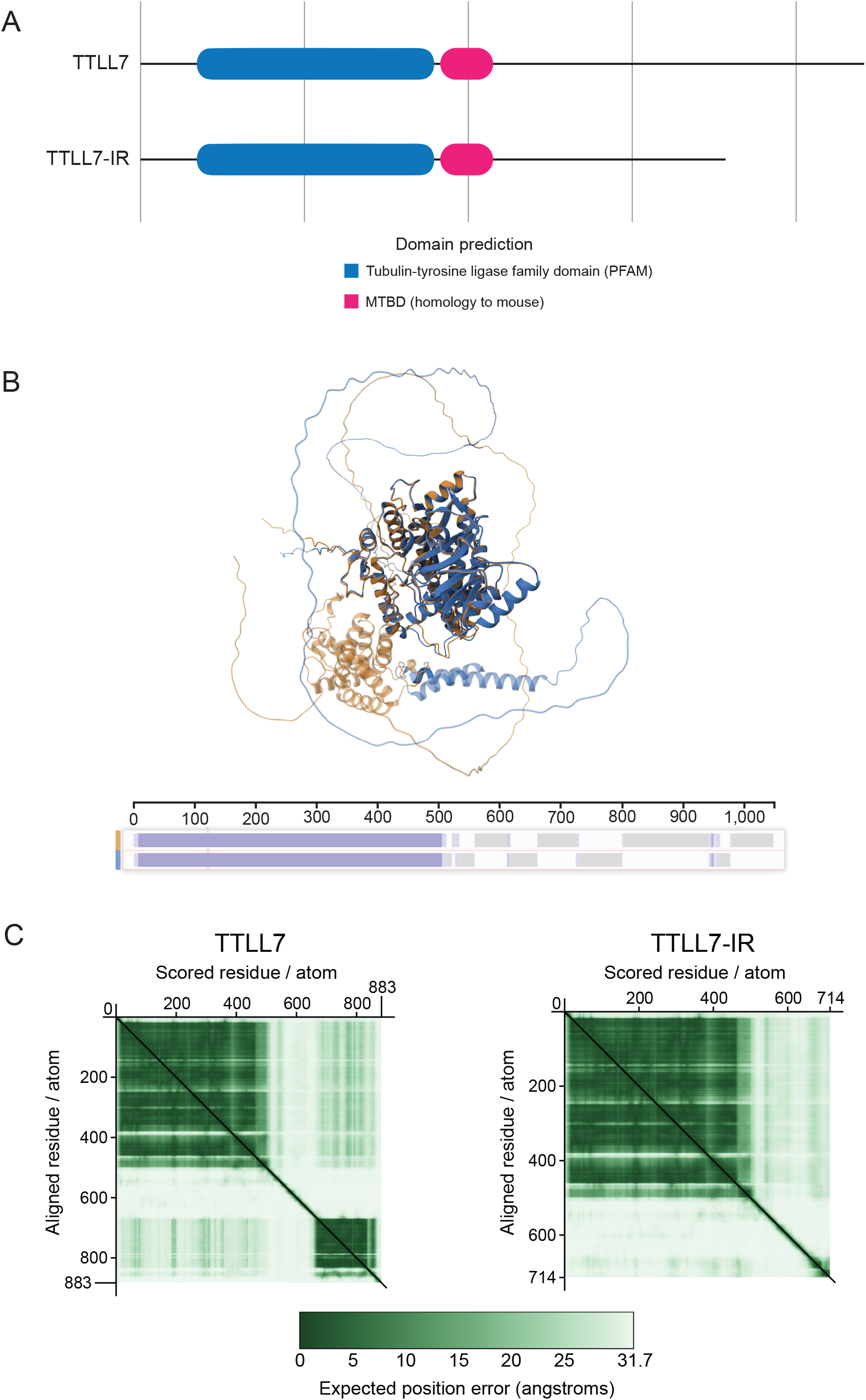
Protein prediction and alignment of canonical and intron-retained *Ttll7*. **A)** Functional domain prediction of the protein sequence resulting from the translation of mRNAs of canonical *Ttll7* (above), and *Ttll7* retention of intron 16 (*Ttll7-IR*) (below). Vertical grey lines indicate 200 amino acids. MTBD is microtubule binding domain. **B)** 3D Protein alignment of the predicted tertiary structure of TTLL7 (yellow) and TTLL7-IR (blue). Transparency indicates no alignment. Amino acid sequence alignment for TTLL7 and TTLL7-IR is shown below with total amino acid sequence number on the x-axis. Purple indicates aligned residues; grey indicates no alignment. **C)** Aligned residue score for predicted protein structure of TTLL7 (left) and TTLL7-IR (right). Protein structures for B and C were generated with AlphaFold Server.

### Intron retention in histone modifying genes

In studies of the central bearded dragon and the two TSD species, the genes *Jarid2* and *Kdm6b* were found to have differentially retained introns between sexes, and specifically in sex-reversed ZZf in the bearded dragon compared to the canonical sexes (ZW females and ZZ males) (Deveson et al. 2017; Whiteley et al. 2022). We investigated whether intron retention was also present in these genes in *B. duperreyi*. Intron retention in both *Kdm6b* and *Jarid2* was present in all sex conditions in *B. duperreyi*. Intron 15 in *Jarid2* was retained completely, with reads covering both exon boundaries. The IR ratio, generated with IRFinder, for intron 15 was also substantially higher than all other introns in *Jarid2* (Figure 4). The same was true for *Kdm6b*, which had IR of intron 22, including reads covering both exon boundaries and a much higher IR ratio than all other introns (Figure 4).

**Figure 4.**
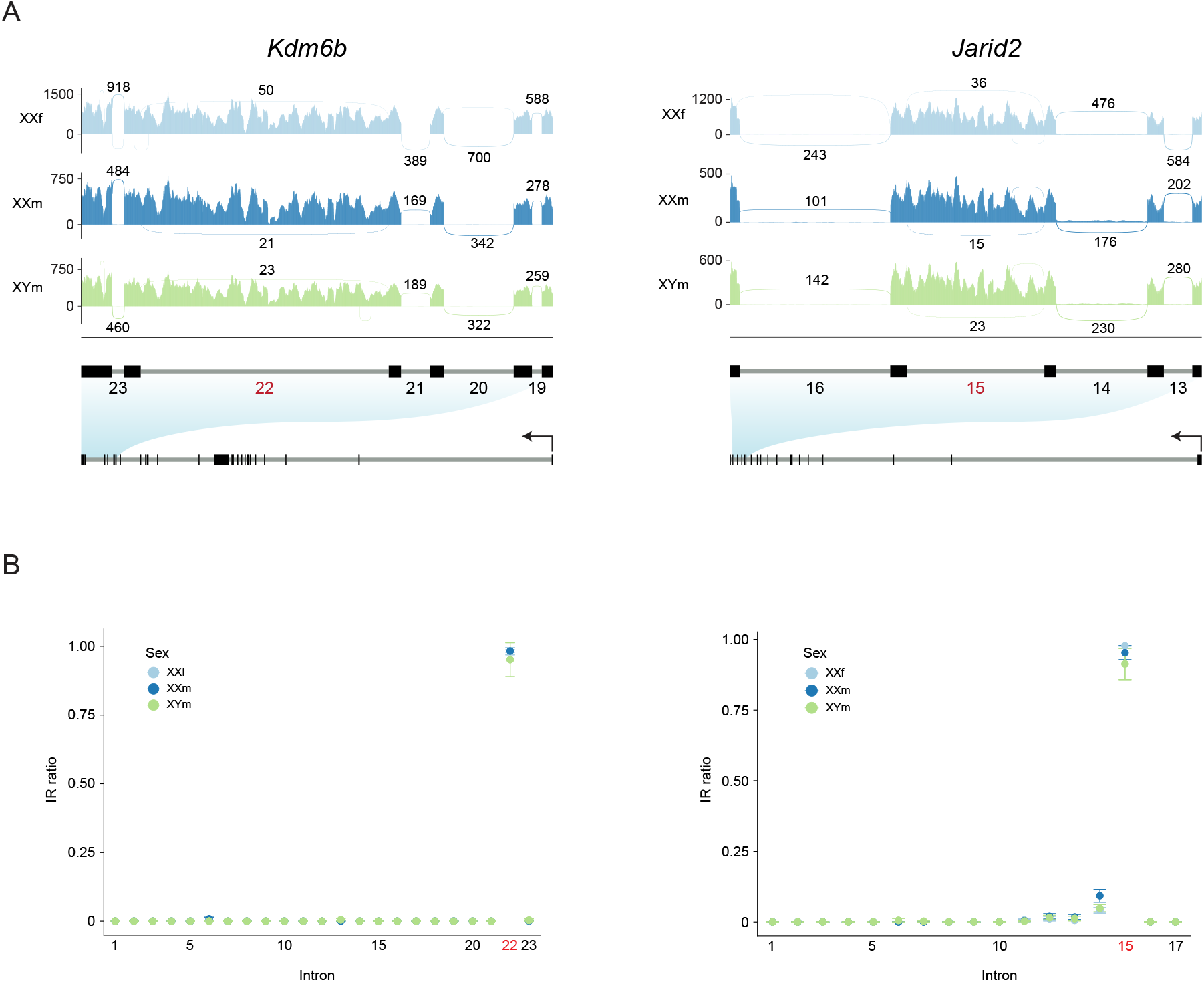
Intron retention occurs in *Kdm6b* and *Jarid2* in all sexes of *B. duperreyi*. **A)** RNA-seq coverage of *Kdm6b* exons 19-24 (left) and *Jarid2* exons 14-17 (right). Tracks show the mean read coverage for each sex condition, XX female (light blue, n = 3), XX male (dark blue, n = 2) and XY male (green, n = 3). The mean number of reads spanning exon junctions are shown as lines spanning the intron above or below the track for each condition. Exon positions are shown below the sashimi plot, including a full gene context. **B)** Intron retention ratio for each intron of *Kdm6b* (left) and *Jarid2* (right) as reported by IRFinder. Points represent the mean IR ratio for each sex condition, XX female (light blue, n = 3), XX male (dark blue, n = 2) and XY male (green, n = 3) and bars are standard error.

We also analysed datasets of adult canonical *B. duperreyi* XY males and XX females, (brain, liver and gonad) transcriptomes for IR. These tissues all showed intron retention in the same introns in *Jarid2* and *Kdm6b* in both sexes and in all tissues (Figure S1). Intron retention in *Jarid2* and *Kdm6b* is therefore not restricted to brain tissue, nor is it exclusive to an early life stage in *B. duperreyi*.

## Discussion

In this study we demonstrate that the expression profile of *B. duperreyi* in brains of sex-reversed XX males is more similar to XX females than XY males. We also identify that differential intron retention is present in XXm male brains in the gene *Ttll7* compared to both canonical sexes. The genes *Kdm6b* and *Jarid2*, which are implicated in another lizard with sex reversal also show intron retention in *B. duperreyi*; however, it was present in all sex conditions.

Our findings that the brain transcriptome of sex-reversed XX males in *B. duperreyi* is more like that of XX females than XY males is concordant with our recent finding that X chromosomes dosage in XX males is the same as XX females and distinct from XY males (Hanrahan et al. 2026). XY males showed partial dosage compensation of the X chromosome of 65-70 % when compared to XX females, whereas XX males were not different from XX females. Here, gene expression levels in XX males and XX females were more different from XY males with 32 and 35 DEGs respectively, than to each other, with only 4 DEGs (Table S3, S4). Therefore, if the signal causing sex reversal in XX males is causing or caused by changes in gene expression, this must occur at an early stage of development not sampled here, or occur specifically in the gonad, or both.

The small size of *B. duperreyi*, up to 90 mm snout to tail length as adults (Cogger 2014), is prohibitive of RNA extraction and successful library preparation from small tissues such as gonad from developing embryos. As such, transcriptomes from an informative period during development have not been studied. Single cell RNA-seq would be an interesting avenue of further research.

We report one novel intron retention event in *B. duperreyi* in the gene *Ttll7*. Intron 16 is retained in sex-reversed XX males at a rate of 36% and is not retained in XX females or XY males (Figure 2). TTLL7 is beta-tubulin polyglutamylase with a role in neurite growth, and has been shown to have brain specific expression in mammals (Ikegami et al. 2006). The intron retained variant of *Ttll7*, if translated, would result in a truncated protein (Figure 3). The effect the truncated protein would have on the brain of *B. duperreyi* is unclear. However, studies of mammalian *Ttll7* have shown that in cells overexpressing a truncated *Ttll7*, similar to that predicted to occur from IR, did not reduce glutamylation activity. In fact, activity of truncated TTLL7 has increased glutamylation activity compared to the full-length protein (van Dijk et al. 2007).

Our finding that *Ttll7* intron retention occurs in sex-reversed XX males in *B. duperreyi* may indicate a change in the physiology of XX male brains. In the central bearded dragon, a lizard which also shows sex reversal at temperature extremes during development, sex-reversed ZZ females show altered behaviours compared to canonical ZW females. These include boldness and activity levels more like genetic ZZ males (Li et al. 2016). Currently there are no studies of the behaviour of sex-reversed individuals in *B. duperreyi* to suggest whether it may be discordant with their phenotypic sex. Further study of the molecular biology of brains and general behaviour in sex-reversed *B. duperreyi* may reveal key insights into physiological changes that result from sex reversal.

### *Kdm6b* and *Jarid2*

The histone modifying genes *Kdm6b* and *Jarid2*, and intron retention specifically, have been implicated in the mechanism of sex reversal in the central bearded dragon (Deveson et al. 2017; Whiteley et al. 2022). They have also posited involvement in the temperature signal from TSD species such as *A. mississippiensis* and *T. scripta* (Czerwinski et al. 2016; Yatsu et al. 2016; Ge et al. 2018). Our observation that *B. duperreyi* has intron retention in both *Jarid2* and *Kdm6b* in all sexes (Figure 4), contrasts that seen in sex reversal of the bearded dragon, which shows IR in *Jarid2* and *Kdm6b* only in sex-reversed ZZ females in adults (Deveson et al. 2017), and in both canonical sexes (ZZm males and ZWf females, 28°C low temperature incubations) and not in ZZf females in developing gonads (Whiteley et al. 2022).

*B. duperreyi* sex reversal occurs at low incubation temperatures, which contrasts the high temperatures required for sex reversal in bearded dragon (Quinn et al. 2007). In *B. duperreyi* incubation temperatures of 23°C result in balanced sex ratios (Radder et al. 2008), and ∼16°C results in sex reversal. These temperatures do not approach those required (∼32-36°C) for sex reversal in the bearded dragon (Quinn et al. 2007).

As such, similar dynamics may not be at play in *B. duperreyi* as seen in bearded dragon where high temperatures will trigger the splicing of the retained intron in *Jarid2* and *Kdm6b*. Possibly these genes are not directly involved in sex determination in *B. duperreyi* and a different mechanism for sex reversal is the trigger at low temperatures. Several genes are known to have altered activity at colder temperatures, such as the gene Cold-inducible RNA-binding protein (*Cirbp*) which has been proposed as a regulator of TSD in the snapping turtle (*Chelydra serpentina*) due to allelic variants being linked to thermosensitivity (Schroeder et al. 2016). *Cirbp* is also differentially expressed during development in other TSD species; the Chinese alligator (*Alligator sinensis*) (Lin et al. 2018), the painted turtle (*Chrysemys picta*) and the spiny softshell turtle (*Apalone spinifera*) (Radhakrishnan et al. 2017). It was not identified as differentially expressed or spliced in our analysis of *B. duperreyi*. However, it would be an interesting target for future work on *B. duperreyi* focussing on the critical thermosensitive window during development due to its implication in TSD species.

## Materials and Methods

### Sample collection and sexing

Full sample collection methods are described in Hanrahan et al. (2026), egg collection is described in Dissanayake et al. (2021b) and a description of the collection site can be found in Dissanayake et al. (2022). In brief, eggs of alpine *B. duperreyi* were collected from nests in the field after ensuring 90% of the development period had passed. They were then transported to the University of Canberra, where they were buried in moist vermiculite with a ratio of 4 parts water to 5 parts vermiculite by weight, and placed in incubators (LabWit, ZXSDR1090) that maintained 23°C until hatching. Phenotypic sex was determined by hemipene eversion at 7 days old and hemipene transillumination at 5 weeks. Genotypic sex was determined using polymerase chain reaction (PCR)-based molecular sex tests on DNA extracted from tail snips. Sex-reversal status was determined using PCR and Y-specific markers, allowing XYm males to be identified and differentiated from XXm males. The lizards were euthanised via intraperitoneal injection of sodium pentobarbitone (100–150 μg/g body weight) as approved by the University of Canberra Animal Ethics Committee (AEC 17–26).

### RNA extraction and sequencing

RNA extraction and sequencing methods are available in Hanrahan et al. (2026) for the brain tissue of the three XY males, three XX females, and two XX male (XXm) *B. duperreyi* hatchlings used here. This resulted in the generation of seventy-five bp single-ended reads on the Illumina NextSeq 500 platform at the Ramaciotti Centre for Genomics (UNSW, Sydney, Australia).

### Differential gene expression

Raw read quality of RNA sequencing was assessed using FastQC v0.11.9 (Andrews 2010) and trimmed accordingly with trimmomatic v0.38 (Bolger et al. 2014). Trimmed reads were aligned to the *B. duperreyi* genome (GenBank accession: GCA_041722995.2) with the subread-align function in subread v2.0.6 (Liao et al. 2013) and summarised with featureCounts. Differential gene expression was determined using the edgeR package v4.6.3 in R v4.5.1 (Chen et al. 2025). Counts were normalised to library size and filtered using the calcNormFactors and filterByExpr functions respectively with default arguments. DEGs were calculated using the quasi-likelihood (QL) pipeline as outlined in the edgeR User’s Guide.

### Intron retention

To quantify intron retention, reads were trimmed as above and aligned to the *B. duperreyi* genome (GenBank accession: GCA_041722995.2) using STAR v2.7.9a as part of the nf-core/rnasplice pipeline (Dobin et al. 2013; Ewels et al. 2020). IRFinder v1.3.1 was then used to quantify intron retention (Middleton et al. 2017). DEseq2 v 1.46.0 (Love et al. 2014) was used to test for differentially retained introns in a pairwise fashion between all sex conditions (XYm male, XXf female and XXm male) in R v4.5.1 as outlined in the IRFinder manual (available at https://github.com/williamritchie/IRFinder/wiki). The analysis was performed using a generalised linear model (GLM) approach as recommended for tests with samples >=3 replicates in some conditions. The thresholds differentially retained introns were a change of > 10% and adjusted p value < 0.05. The Benjamini-Hochberg method was used to adjust p values. In total 180,950 introns were tested.

### Protein prediction and alignment

The TTLL7 protein sequence was predicted based on translation of the *B. duperreyi* assembly and annotation. Functional domain prediction was performed using InterProScan v5.76-107.0 and alignment from functional domains in mouse sourced from UniProt (ID: A4Q9F0) using blastp sequence alignment (Camacho et al. 2009; Jones et al. 2014; UniProt 2025). The 3D protein structure of TTLL7 was predicted using AlphaFold Server for full length and intron retained transcripts of *Ttll7* (Abramson et al. 2024). The highest confidence predictions for each of TTLL7 and TTLL7-IR were then aligned with the PDB structural alignment tool with TM-align as the alignment method (Bittrich et al. 2024).

## Declarations

## Data access

All raw sequencing data used in this study is available from the NCBI BioProject database (https://www.ncbi.nlm.nih.gov/bioproject/) under accession number PRJNA980841. The genome assembly used in this study is available from the NCBI Genome database under GenBank accession GCA_041722995.2.

## Acknowledgments

N/A

## Ethics approval

The animals involved in this study were collected from nests in the field under permits issued by the Australian state government: ACT permit numbers (LT201826, LT2017956) and NSW government (SL102002). The collection was carried out in accordance with their respective guidance and protocols, as well as the University of Canberra Animal Ethics Committee (Approval Number AEC 17–26). All husbandry practices complied with the Australian Code for the Care and Use of Animals for Scientific Purposes, 8th edition (2013), particularly sections 3.2.13–3.2.2.

## Author contributions

B.J.H wrote the original manuscript and incorporated edits based on feedback. B.J.H, P.D.W and A.G designed the study. D.S.B.D collected and harvested samples. B.J.H and J.K.C conducted extraction experiments to generate data. B.J.H conducted data analysis and visualisation. N.C.L provided insight into data interpretation and writing. All authors reviewed and approved the submitted manuscript.

## Supplementary figure legends

**Figure S1. Intron retention occurs in *Kdm6b* and *Jarid2* in adult tissues of XY male and XX female *B. duperreyi***. RNA-seq coverage of **A)** *Jarid2* from exon 14-17 and **C)** *Kdm6b* from exon 19-24. Tracks show the mean read coverage for XX female (light blue) and XY male (green) from brain (n=3), liver (n=1) and gonad (n=1). The mean number of reads spanning exon junctions are shown as lines spanning the intron above or below the track for each tissue. Exon and intron positions are shown below each sashimi plot, introns are numbered. Intron retention ratio is plotted for each intron of **B)** *Jarid2* and **D)** *Kdm6b* as reported by IRFinder. Points represent the IR ratio for each tissue except for brain in which the points are the mean value and bars are standard error.

## Supplementary tables

**Table S1. List of differentially expressed genes between the three sex conditions: XYm, XXf and XXm**.

**Table S2. Top ten blast hits for the coding sequence of the gene *g11335***

**Table S3. Gene Ontology results of differentially expressed genes between XX females and XY males**.

**Table S4. Gene Ontology results of differentially expressed genes between XY males and sex-reversed XX males**.

**Table S5. Results of differential intron retention analysis between the three sex conditions: XYm, XXf and XXm**. A single gene, *Ttll7*, shows intron retention specific to sex-reversed XXm which is not present in XXf or XYm.

## References

Abramson J, Adler J, Dunger J, Evans R, Green T, Pritzel A, Ronneberger O, Willmore L, Ballard AJ, Bambrick J, Bodenstein SW, Evans DA, Hung CC, O’Neill M, Reiman D, Tunyasuvunakool K, Wu Z, Zemgulyte A, Arvaniti E, Beattie C, Bertolli O, Bridgland A, Cherepanov A, Congreve M, Cowen-Rivers AI, Cowie A, Figurnov M, Fuchs FB, Gladman H, Jain R, Khan YA, Low CMR, Perlin K, Potapenko A, Savy P, Singh S, Stecula A, Thillaisundaram A, Tong C, Yakneen S, Zhong ED, Zielinski M, Zidek A, Bapst V, Kohli P, Jaderberg M, Hassabis D, Jumper JM. 2024. Accurate structure prediction of biomolecular interactions with AlphaFold 3. Nature 630: 493–500 doi:10.1038/s41586-024-07487-w.

Andrews S. 2010. FastQC: a quality control tool for high throughput sequence data. Available online at: https://www.bioinformatics.babraham.ac.uk/projects/fastqc.

Bittrich S, Segura J, Duarte JM, BURLey SK, Rose Y. 2024. RCSB protein Data Bank: exploring protein 3D similarities via comprehensive structural alignments. Bioinformatics 40 doi:10.1093/bioinformatics/btae370.

Bolger AM, Lohse M, Usadel B. 2014. Trimmomatic: a flexible trimmer for Illumina sequence data. Bioinform 30: 2114–2120 doi:10.1093/bioinformatics/btu170.

Bull JJ. 1980. Sex Determination in Reptiles. Q Rev Biol 55: 3–21 doi:10.1086/411613.

Bull JJ. 1983. Evolution of sex determining mechanisms. Benjamin/Cummings Pub. Co., Menlo Park, CA.

Camacho C, Coulouris G, Avagyan V, Ma N, Papadopoulos J, Bealer K, Madden TL. 2009. BLAST+: architecture and applications. BMC Bioinformatics 10: 421 doi:10.1186/1471-2105-10-421.

Chen Y, Chen L, Lun ATL, Baldoni PL, Smyth GK. 2025. edgeR v4: powerful differential analysis of sequencing data with expanded functionality and improved support for small counts and larger datasets. Nucleic Acids Res 53 doi: 10.1093/nar/gkaf018.

Cogger H. 2014. Reptiles and Amphibians of Australia. CSIRO PUBLISHING, Melbourne, VIC.

Czerwinski M, Natarajan A, Barske L, Looger LL, Capel B. 2016. A timecourse analysis of systemic and gonadal effects of temperature on sexual development of the red-eared slider turtle Trachemys scripta elegans. Dev Biol 420: 166–177 doi:10.1016/j.ydbio.2016.09.018.

Deveson IW, Holleley CE, Blackburn J, Marshall Graves JA, Mattick JS, Waters PD, Georges A. 2017. Differential intron retention in Jumonji chromatin modifier genes is implicated in reptile temperature-dependent sex determination. Sci Adv 3: e1700731 doi:10.1126/sciadv.1700731.

Dissanayake DSB, Holleley CE, Deakin JE, Georges A. 2021a. High elevation increases the risk of Y chromosome loss in Alpine skink populations with sex reversal. Heredity (Edinb) 126: 805–816 doi:10.1038/s41437-021-00406-z.

Dissanayake DSB, Holleley CE, Georges A. 2021b. Effects of natural nest temperatures on sex reversal and sex ratios in an Australian alpine skink. Sci Rep 11: 20093 doi:10.1038/s41598-021-99702-1.

Dissanayake DSB, Holleley CE, Sumner J, Melville J, Georges A. 2022. Lineage diversity within a widespread endemic Australian skink to better inform conservation in response to regional-scale disturbance. Ecol Evol 12: e8627 doi:10.1002/ece3.8627.

Dobin A, Davis CA, Schlesinger F, Drenkow J, Zaleski C, Jha S, Batut P, Chaisson M, Gingeras TR. 2013. STAR: ultrafast universal RNA-seq aligner. Bioinformatics 29: 15–21 doi:10.1093/bioinformatics/bts635.

Donnellan SC. 1985. The evolution of sex chromosomes in scincid lizards. Doctoral Thesis, Macquarie University, Maquarie Park, NSW.

Ewels PA, Peltzer A, Fillinger S, Patel H, Alneberg J, Wilm A, Garcia MU, Di Tommaso P, Nahnsen S. 2020. The nf-core framework for community-curated bioinformatics pipelines. Nat Biotechnol 38: 276–278 doi:10.1038/s41587-020-0439-x.

Ge C, Ye J, Weber C, Sun W, Zhang H, Zhou Y, Cai C, Qian G, Capel B. 2018. The histone demethylase KDM6B regulates temperature-dependent sex determination in a turtle species. Science 360: 645–648 doi:10.1126/science.aap8328.

Hanrahan BJ, Chang JK, Milton AM, Lister NC, Dissanayake DSB, Hammond JM, Reis ALM, Deveson IW, Ruiz-Herrera A, Patel HR, Graves JAM, Georges A, Waters PD. 2026. Sex chromosome dosage compensation in a sex reversing skink is not influenced by sexual phenotype. BMC Genomics 27: 72 doi:10.1186/s12864-025-12217-1.

Holleley CE, O’Meally D, Sarre SD, Marshall Graves JA, Ezaz T, Matsubara K, Azad B, Zhang X, Georges A. 2015. Sex reversal triggers the rapid transition from genetic to temperature-dependent sex. Nature 523: 79–82 doi:10.1038/nature14574.

Ikegami K, Mukai M, Tsuchida J, Heier RL, Macgregor GR, Setou M. 2006. TTLL7 is a mammalian beta-tubulin polyglutamylase required for growth of MAP2-positive neurites. J Biol Chem 281: 30707–30716 doi:10.1074/jbc.M603984200.

Jones P, Binns D, Chang HY, Fraser M, Li W, McAnulla C, McWilliam H, Maslen J, Mitchell A, Nuka G, Pesseat S, Quinn AF, Sangrador-Vegas A, Scheremetjew M, Yong SY, Lopez R, Hunter S. 2014. InterProScan 5: genome-scale protein function classification. Bioinformatics 30: 1236–1240 doi:10.1093/bioinformatics/btu031.

Li H, Holleley CE, Elphick M, Georges A, Shine R. 2016. The behavioural consequences of sex reversal in dragons. Proceedings of the Royal Society B: Biological Sciences 283 doi:10.1098/rspb.2016.0217.

Liao Y, Smyth GK, Shi W. 2013. The Subread aligner: fast, accurate and scalable read mapping by seed-and-vote. Nucleic Acids Res 41: e108 doi:10.1093/nar/gkt214.

Lin JQ, Zhou Q, Yang HQ, Fang LM, Tang KY, Sun L, Wan QH, Fang SG. 2018. Molecular mechanism of temperature-dependent sex determination and differentiation in Chinese alligator revealed by developmental transcriptome profiling. Sci Bull (Beijing) 63: 209–212 doi:10.1016/j.scib.2018.01.004.

Love MI, Huber W, Anders S. 2014. Moderated estimation of fold change and dispersion for RNA-seq data with DESeq2. Genome Biol 15: 550 doi:10.1186/s13059-014-0550-8.

Matsubara K, O’Meally D, Azad B, Georges A, Sarre SD, Graves JA, Matsuda Y, Ezaz T. 2016. Amplification of microsatellite repeat motifs is associated with the evolutionary differentiation and heterochromatinization of sex chromosomes in Sauropsida. Chromosoma 125: 111–123 doi:10.1007/s00412-015-0531-z.

Middleton R, Gao D, Thomas A, Singh B, Au A, Wong JJ, Bomane A, Cosson B, Eyras E, Rasko JE, Ritchie W. 2017. IRFinder: assessing the impact of intron retention on mammalian gene expression. Genome Biol 18: 51 doi:10.1186/s13059-017-1184-4.

Quinn AE, Georges A, Sarre SD, Guarino F, Ezaz T, Graves JA. 2007. Temperature sex reversal implies sex gene dosage in a reptile. Science 316: 411 doi:10.1126/science.1135925.

Quinn AE, Radder RS, Sarre SD, Georges A, Ezaz T, Shine R. 2009. Isolation and development of a molecular sex marker for Bassiana duperreyi, a lizard with XX/XY sex chromosomes and temperature-induced sex reversal. Mol Genet Genomics 281: 665–672 doi:10.1007/s00438-009-0437-7.

Radder RS, Quinn AE, Georges A, Sarre SD, Shine R. 2008. Genetic evidence for co-occurrence of chromosomal and thermal sex-determining systems in a lizard. Biol Lett 4: 176–178 doi:10.1098/rsbl.2007.0583.

Radhakrishnan S, Literman R, Neuwald J, Severin A, Valenzuela N. 2017. Transcriptomic responses to environmental temperature by turtles with temperature-dependent and genotypic sex determination assessed by RNAseq inform the genetic architecture of embryonic gonadal development. PLoS One 12: e0172044 doi:10.1371/journal.pone.0172044.

Schroeder AL, Metzger KJ, Miller A, Rhen T. 2016. A Novel Candidate Gene for Temperature-Dependent Sex Determination in the Common Snapping Turtle. Genetics 203: 557–571 doi:10.1534/genetics.115.182840.

Shine R, Elphick MJ, Donnellan S. 2002. Co-occurrence of multiple, supposedly incompatible modes of sex determination in a lizard population. Ecol Lett 5: 486–489 doi:10.1046/j.1461-0248.2002.00351.x.

UniProt C. 2025. UniProt: the Universal Protein Knowledgebase in 2025. Nucleic Acids Res 53: D609–D617 doi:10.1093/nar/gkae1010.

van Dijk J, Rogowski K, Miro J, Lacroix B, Edde B, Janke C. 2007. A targeted multienzyme mechanism for selective microtubule polyglutamylation. Mol Cell 26: 437–448 doi:10.1016/j.molcel.2007.04.012.

Whiteley SL, Wagner S, Holleley CE, Deveson IW, Marshall Graves JA, Georges A. 2022. Truncated jarid2 and kdm6b transcripts are associated with temperature-induced sex reversal during development in a dragon lizard. Sci Adv 8: eabk0275 doi:10.1126/sciadv.abk0275.

Yatsu R, Miyagawa S, Kohno S, Parrott BB, Yamaguchi K, Ogino Y, Miyakawa H, Lowers RH, Shigenobu S, Guillette LJ, Jr., Iguchi T. 2016. RNA-seq analysis of the gonadal transcriptome during Alligator mississippiensis temperature-dependent sex determination and differentiation. BMC Genomics 17: 77 doi:10.1186/s12864-016-2396-9.

