## Supplementary figures and images for "Characterising differential gene expression and alternative splicing in a sex reversing skink, *Bassiana duperreyi*"

### Figure S1

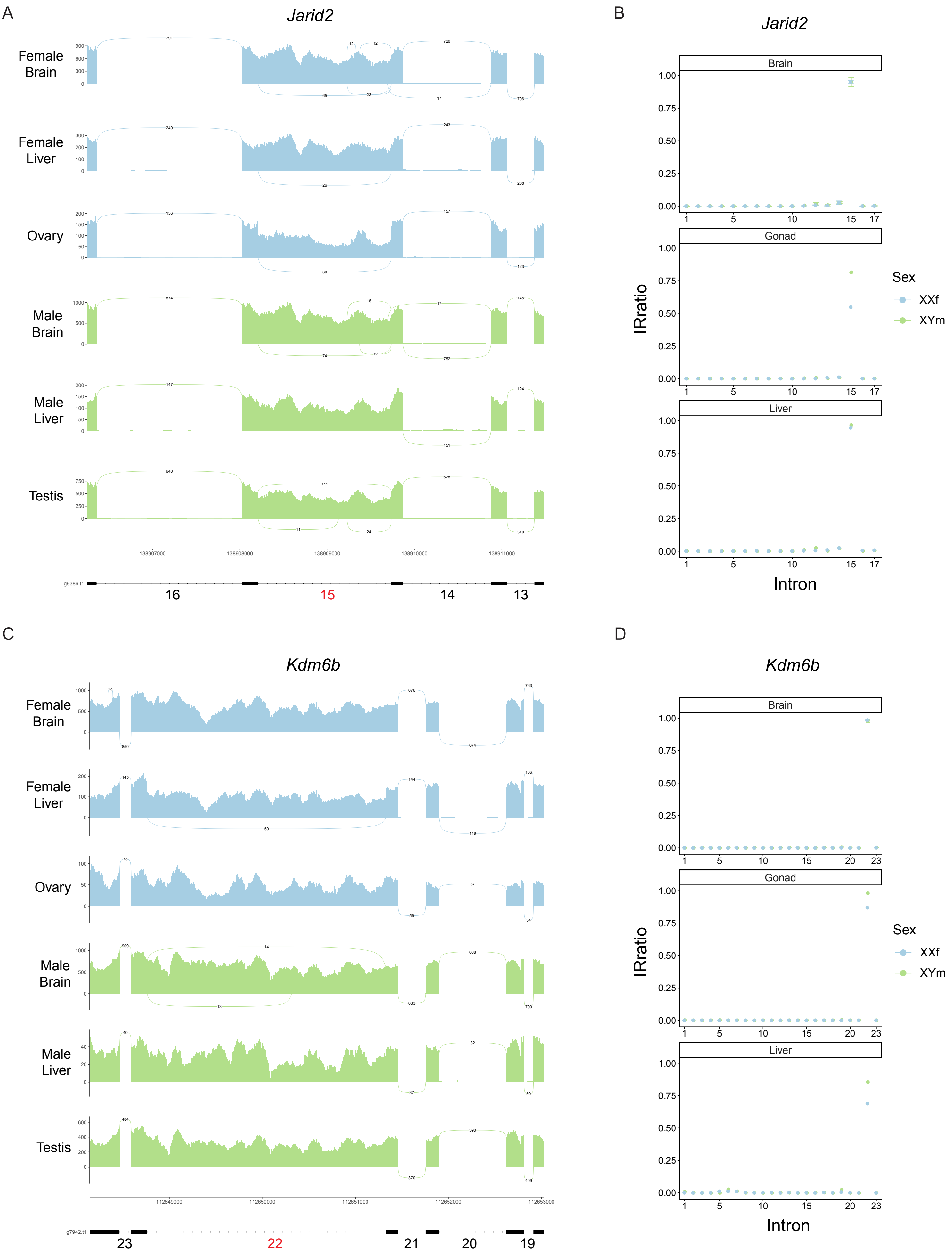
